# Using random forests to uncover the predictive power of distance–varying cell interactions in tumor microenvironments

**DOI:** 10.1101/2023.07.18.549619

**Authors:** Jeremy VanderDoes, Claire Marceaux, Kenta Yokote, Marie-Liesse Asselin-Labat, Gregory Rice, Jack D. Hywood

**Affiliations:** Department of Statistics and Actuarial Science, University of Waterloo, Waterloo, Canada; Personalised Oncology Division, The Walter and Eliza Hall Institute of Medical Research, Parkville, Australia; Department of Medical Biology, The University of Melbourne, Parkville, Australia; Department of Anatomical Pathology, Royal Melbourne Hospital, Melbourne, Australia

## Abstract

Tumor microenvironments (TMEs) contain vast amounts of information on patient’s cancer through their cellular composition and the spatial distribution of tumor cells and immune cell populations. Exploring variations in TMEs among patients and cancer types, as well as determining the extent to which this information can predict variables such as patient survival or treatment success with emerging immunotherapies, is of great interest. Moreover, in the face of a large number of potential spatial cell interactions to consider, we often wish to identify specific interactions that are useful in making such predictions. We present an approach to achieve these goals based on summarizing spatial relationships in the TME using spatial *K* functions, and then applying functional data analysis and random forest models to both predict outcomes of interest and identify important spatial relationships. This approach is shown to be effective in simulation experiments. We further used the proposed approach to interrogate two real data sets of Multiplexed Ion Beam Images of TMEs in triple negative breast cancer and lung cancer patients. The methods proposed are publicly available in a companion `R` package `funkycells`.

**Author summary:** Spatial data on the tumor microenvironment (TME) are becoming more prevalent. Existing methods to interrogate such data often have several deficiencies: (1) they rely on estimating the spatial relationships among cells by examining simple counts of cells within a *single* radius, (2) they do not come with ways to evaluate the statistical significance of any findings, or (3) they consider multiple individual interactions resulting in overly optimistic estimates of interaction importances. Our approach, which leverages techniques in spatial statistics and uses a benchmark ensemble machine learning method addresses (1), since the *K* functions used encode the relative densities of cells over all radii up to a user-selected maximum radius, and (2) we have developed a custom approach based on permutation and cross-validation to evaluate the statistical significance of any findings of significant spatial interactions in the TME, (3) over potentially multiple interactions. Our approach is also freely available with an `R` implementation called `funkycells`. In the analysis of two real data sets, we have seen that the method performs well, and gives the expected results. We think this will be a robust tool to add to the toolbox for researchers looking to interrogate, what can be sometimes unwieldy, TME data.

## Introduction

Recent advances in cancer treatment, such as immune checkpoint inhibition and other cancer immunotherapies, have sparked a growing interest in studying the cellular composition and spatial organization of the tumor microenvironment (TME). The latest innovations in imaging technologies allow for single cell resolution of specific proteins, facilitating in-depth study of the spatial arrangement of cell types within the TME. A wide variety of technologies are available for this purpose, each with different benefits and trade-offs [1–7]. For a review of the available technologies see [8].

In comparing TME data, different spatial relationships between cell types, e.g. between tumor cells and specific immune cell populations, and/or individual proteins, often appear predictive of patient outcomes and may guide therapeutic interventions; see for example [9]. Comparisons between cancer subtypes, e.g. hormone-positive versus hormone-negative breast cancers, or lung squamous cell carcinoma vs lung adenocarcinoma, may provide novel insight into tumor biology and guide the development of treatments. A further goal is to identify specific spatial relationships observed in particular patient’s tumor that are useful in predicting an outcome, such as patient survival or response to therapy. Recent results demonstrate that TME data can be used for such prediction in a variety of tumor types [10, 11].

We consider such prediction problems for data sets generated from tumors imaged with Multiplexed Ion Beam Imaging (MIBI) by means of the MIBIscope in this paper. The MIBIscope uses ion-beam ablation and time-of-flight mass spectrometry to detect up to 40 protein markers on formalin-fixed, paraffin-embedded (FFPE) tissue. Thus, it provides deep data on cell characteristics and their localisation at a single-cell resolution of around 250-400nm [12, 13]. Data collected on the TME, using e.g. the MIBIscope, can be considered as marked spatial point patterns [14–16]. The cell locations can be considered as points within the pattern, with cell phenotypes and/or protein markers giving the “marks”. An example of such a point process generated from a tumor imaged using MIBIscope is shown in the left hand panel of Fig 1.

**Fig 1.**
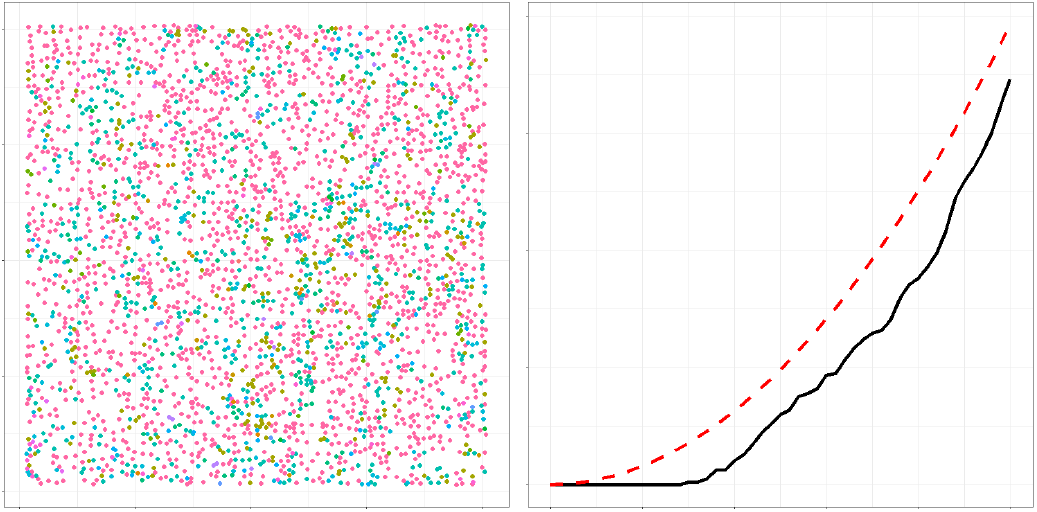
Sample point pattern and *K* function. A point pattern produced for a tumor imaged using MIBIscope from a triple negative breast cancer patient with multiple identified phenotypes (left) and the associated cross K function (black) for two cell types in the image: tumor and monocytes/neutrophils (right). In the tumour, the *x*- and *y*-axes represent the spatial dimensions while in the *K* function,the *x*-axis indicates the radius of interest and the *y*-axis gives the value of the *K* function. The point pattern contains 15 uniquely colored phenotypes, including tumor (red), NK (purple), and monocytes/neutrophils (cyan). The cross-over *K* function has a red reference line gives *πr*^2^ (red), associated with complete spatial randomness.

Methods developed to this point to study cellular interactions in the TME have exploited cell neighborhood analysis in which the spatial relationship between a cell of interest and its neighboring cells can reveal particular cell-cell interactions associated with a disease state or changes associated with response to therapy; see e.g. [17].

Pairwise cell-to-cell distance calculations over iterations of randomized permutations has also been used to identify relevant cell-cell interactions [18]. However, the substantial number of cell types present in the TME leads to a very large number of potential pairwise interactions creating a major challenge in finding interactions that may be meaningful and statistically significant in predicting outcomes of interest. There have been many investigations into applying spatial statistics methods to similar biological data sets, e.g. [19], [20], and [21].

A common method of analysing spatial point patterns, such as those that arise in TME imaging, is to consider Ripley’s *K* function [22–24]. The *K* function describes the distribution of inter-point distances in a given point pattern, giving an indication as to whether points in the pattern (e.g. cells) are clustered or dispersed with respect to one another. The *K* function, along with other summary functions from spatial statistics, has previously been employed in the analysis of the TME [25–31]. An example of a *K* function showing the relative distribution of a specific immune cell type around tumor cells within a MIBIscope image from a triple negative breast cancer patient is shown in the right hand panel of Fig 1.

In this paper, we present a general framework for analyzing and identifying useful spatial relationships in the TME through predicting an outcome of interest. The method we propose uses a novel combination of spatial statistics and functional data analysis, in conjunction with methods in ensemble machine learning. The application of functional data analysis to spatial statistics is a recent development [31–35].

Our approach begins by producing *K* functions for the different cell-cell, (or alternatively marker-marker), interactions within images. After performing dimension reduction using functional principal components analysis [36], these data are combined with non-functional patient meta-data, such as age or sex, and a modified random forest model is used to predict the patient outcome. Motivated by [37], in order to evaluate the predictive power of the spatial interactions, “knock-off” point patterns that mimic the spatial data in the TME are generated, via permutation, independently of the responses. The importance of specific spatial interactions in predicting the response are evaluated by comparison to the predictive power of the knock-off spatial patterns. This approach overcomes the challenge of distinguishing important spatial interactions among many potential interactions of interest, and for the generation of easy-to-interpret plots showing which interactions are useful in predicting the response at a glance. Moreover, it grants high power for even a relatively few number of cells due to the robustness of the *K* functions, and high power for small sample sizes due to the knock-offs. Small sample sizes are common in rare cancer analyses, but are of increasing interest [19].

In many analyses, a single distance of interest is considered while here we consider a range of distances through *K* functions. [30] provides an approach for detecting differences across multiple images between cell-cell interactions by comparing the integrated difference between the empirical *L* function (a function derived from the *K* function, see [38]) and the *L* function associated with complete spatial randomness. However, this approach independently considers the interactions, while we wish to consider all interactions in a single model. Additionally, a possible drawback of the approach in [30] is that integration over the summary statistic may lose valuable information relating to differences in the shape of functions (i.e. differently shaped *L* functions may still have the same integral).

Some methods also consider only a single images or an equal number of images per patient. However, our approach can be used to analyse data with multiple, possibly differing, numbers of observations per patient. In that sense, it allows for the complete use of the data, rather than removing data.

We apply the proposed methods to two MIBIscope data sets; a data set of triple negative breast cancer (TNBC) patients, and a data set consisting of both lung squamous cell carcinomas (LUSC) and lung adenocarcinomas (LUAD). Regarding the TNBC data, our method was accurately able to identify clustered versus dispersed tumors when compared to [12], and was additionally able to identify important cell spatial interactions in making that determination. Our method also indicated that there did not appear to be measurable differences in the spatial arrangement of tumor and immune cell types, as measured by K functions, between the LUSC and LUAD groups.

Whilst the methodology presented here is motivated by, and applied to, MIBIscope data, it can be applied to similar data generated by other technologies, e.g. OPAL, Phenocycler Fusion, Merscope, Xenium and Cosmx [39–45]. Furthermore, the methodology can easily extend beyond two-dimensions to higher-dimensional images, another area of active research [46].

The rest of the paper is organized as follows. In Materials and methods, we give a detailed description of the data we consider and the methods to analyze them, including sub-sections on how we fit a modified random forest in this setting, and how we evaluate the statistical significance and uncertainty in measuring the variable importance of spatial interactions of cell types as encoded by *K* functions. We also introduce the `R` package `funkycells`, an open-source implementation of our approach in that section. Simulation study details the results of simulation experiments in which we found that the proposed method performed well when applied to synthetic data built to mimic the TNBC data. We report the results when this approach was applied to the TNBC and LUSC vs. LUAD data sets in Applications to MIBIscope data. Some concluding remarks and directions for future work are collected in Discussion.

## Materials and methods

The raw spatial data that we consider take the form of 2-dimensional point patterns, as generated using MIBIscope. We denote the cell spatial data, as

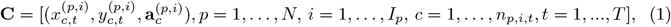

where 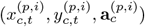 denotes the *x* and *y* coordinates of the *c*^th^ cell of type *t*, of which there are *T* total types, for the *i*^th^ image of the *p*^th^ patient, 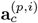 defines cell attributes related to the *c*^th^ cell of type *t* (e.g. phenotype, antigen marker intensities, etc.), and *n*_*p,i,t*_ is the number of cells in image *i* of patient *p* over all types, e.g. phenotypes or proteins. The properties in 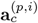 may simply give the cell’s phenotype (and is therefore redundant due to the term *t*), or may be more general, such as a vector describing individual protein expression. For example, the vector could be composed of binary indicators as to whether a protein is expressed or not. For notational clarity, and since we only consider data consisting of cells and their associated phenotype, we drop the **a** term throughout the paper. Fig 1 shows a point pattern for one tumor from the TNBC data indicating several phenotypes, including tumor cells, monocytes and neutrophils (Mono/Neu), and *B* cells [12].

Since the applicability of our method extends beyond this example, we designate several general terms for use throughout the paper. We refer to point patterns such as in Fig 1 as “images”. We interchangeably use the terms cell phenotype and cell type. We also interchangeably use the terms cross-over K function and K function. We assume a single response variable, *Z*_*i*_, for each of the *N* patients (e.g. tumor type, response to therapy, etc.). The set of outcomes for the *N* patients is denoted **Z** = (*Z*_1_, …, *Z*_*N*_). In the real data examples we consider below *Z*_*i*_ is a binary response, e.g. “compartmentalized” versus “mixed” tumors for the TNBC data, or LUSC versus LUAD for the lung cancer data, in which case we can encode the outcomes as taking the values 0 and 1. These methods may easily be adapted for more general class responses, e.g. different types of tumors, or numeric responses, e.g. survival time.

In addition to the spatial data, we assume that we may have access to non-spatial data on the patients. We refer to this data as patient “meta-data”, and we assume that it takes the form **M** = (**m**_1_, …, **m**_*N*_), where each **m**_*i*_ is a vector of patient attributes, for example age or sex.

With both the cell spatial data **C** and meta-data **M**, our goals are to (1) investigate to what extent these data are useful in predicting the outcomes **Z**, and (2) to identify which specific spatial relationships and/or components of the meta-data from the full data set are useful in predicting **Z**. We deem data on a spatial relationship or component of the meta-data “useful” if their importance in predicting the outcome exceeds, to a statistically significant degree, that of similar variables that are known to be unrelated to the outcome. For reference throughout the paper, a high-level schematic of our proposed method is presented in Fig 2.

**Fig 2.**
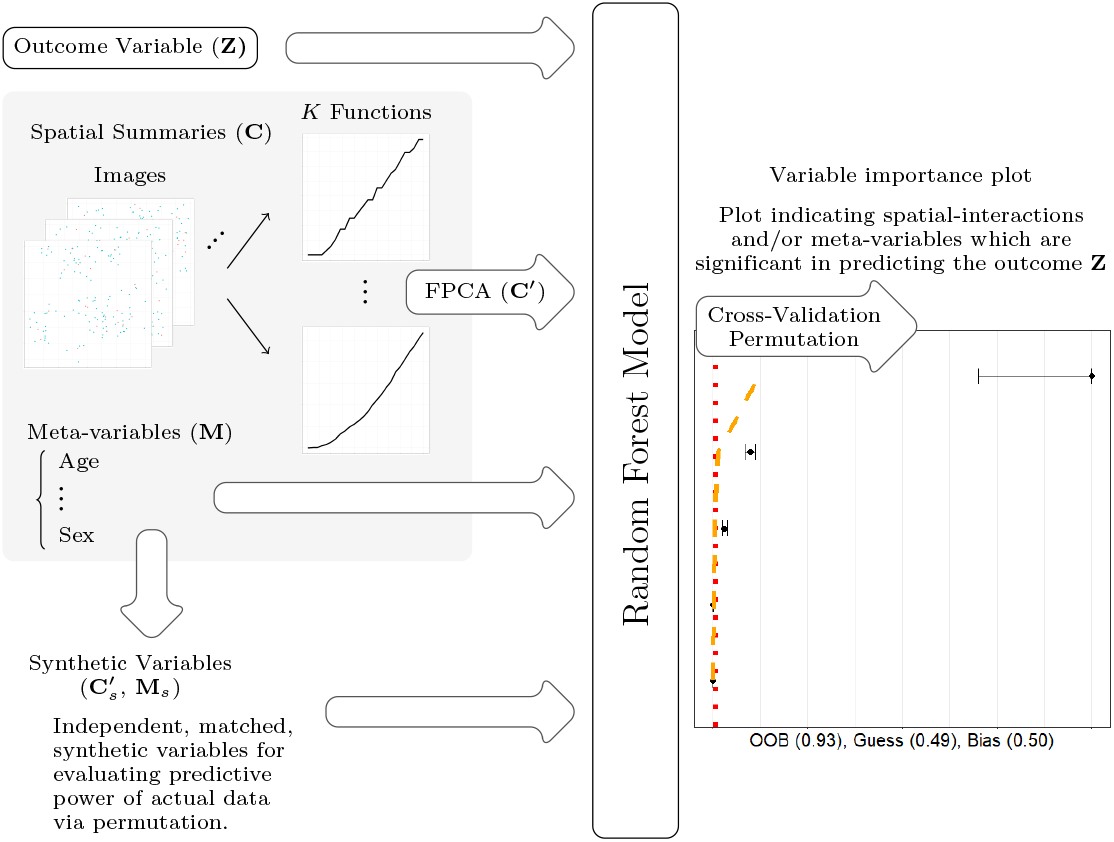
Flow chart of model. When modeling using `funkycells`, there are several major steps: organizing data, generating synthetic data, and modeling using random forests. The spatial data is organized into functional summaries (*K* functions) that are projected into finite dimensions (FPCA) and used with meta-variables to predict the outcome variable. The spatial data and meta-variables are permuted to create synthetic variables with similar properties but independent of the outcome. These synthetic variables are then added to the model, and used to quantify the strength of the relationships between the spatial and meta-data with the response. The model processes the data, employing cross-validation and permutation to return a variable importance plot (with predictive accuracy estimates) indicating spatial interactions and/or meta-variables which are significant in predicting the outcome **Z**.

Towards answering these questions, we build a model of the outcomes **Z** in terms of the image spatial information **C** and meta-data **M**. In doing so, we must address how we incorporate the complex image data into such a model. Motivated by the expectation that patient outcomes are influenced by the relative distribution of various immune cells or protein markers around each other, we begin by computing spatial “cross-over” *K* functions from the image data, which summarize the spatial distribution of cells with respect to one another as in Fig 1.

We provide an open source implementation of our approach in `R` [47] at the site github.com/jrvanderdoes/funkycells, in the package `funkycells`. This implementation also includes the code and data used in the presented simulations and data analyses.

### Summarizing the image data using K-functions

The cross-over *K* function for image *i* of person *p* and cell types *t* and *t*′ is defined as

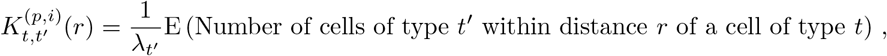

where E denotes mathematical expectation, the radius *r* ranges from 0 ≤ *r* ≤ *R*, with the max radius *R* being a user specified parameter that we discuss below, and *λ*_*t*_*′* gives the density of cells of phenotype *t*′ [14–16].

By examining this function for varying radii, we may infer how cell types are distributed around each other. For example, if cell types are distributed around each other entirely at random, then 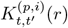 is equal to the area of a circle of radius *r, πr*^2^. Regularity or dispersion of the cells around each other tends to reduce 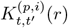 while clustering tends to increase it. An example of a *K* function computed between the tumor and monocytes/neutrophils phenotypes for a given tumor in the TNBC data set is presented in Fig 1, which indicates a degree of dispersion with respect to monocytes and neutrophils cells around tumor cells across *r* values compared to that expected for cells distributed around Tumor cells with complete spatial randomness. Cross-over *K* functions can be used to summarize all two-way interactions between cell phenotypes for a given image.

In practice, estimation is based on an empirical average replacing the expectation. The estimated cross-over *K* function for image *i* of person *p* is given by

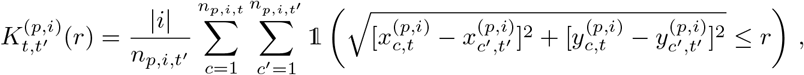

0 ≤ *r* ≤ *R*, where 𝟙 (*A*) takes the value one if the condition *A* is satisfied, and is zero otherwise, and |*i*| indicates the area of the image. When patients have multiple images, we combine their cross-over *K* functions by computing a weighted average,

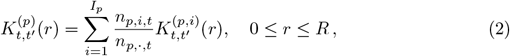

with *I*_*p*_ giving the number of images for a given patient *p*. In other words, the *K* functions from each image are weighted according to the prevalence of the cells of the type under consideration. We note that if there is one image per patient (so that *I*_*p*_ = 1), then 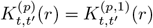, and further that the weights in Eq (2) vanish to zero if the cell types *t* and *t*′ are missing in an image.

It is common when computing such *K* functions to correct them for what are referred to as “edge effects”. Edge effects describe the issue that cells near the edge of an image appear to have fewer cells around them when *r* extends beyond the nearest boundary. One option is toroidal edge corrections as described in [48], which replicates and reflects the image data occurring near the boundary. Another is isotropic edge corrections which weights edge cells, see [49–51]. We examined several methods and found similar results. We use a standard isotropic edge correction in this paper. For a single point *x* in some image *I*, then the isotropic edge correction weight is

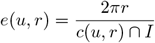

where the denominator is the intersection of the circle *c*(*u, r*) with radius *r* and center *u*. For discussions on approaches to edge corrections see [52] or [53].

In computing these *K* functions for each cell type, we can transform the spatial data **C** into a collection of *T* ^2^ different *K* functions for each patient, 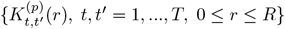. The *K* functions are then treated as functional data objects; see e.g. [36]. Since even moderate values of *T* lead to a large number of *K* functions to consider, user input is often helpful in determining a smaller subset of interactions (and hence *K* functions) of particular interest for analysis.

Although informative, these *K* functions are unwieldy to directly use in a model, and we further transform the functions using the dimension reduction technique of functional principal component analysis (FPCA). FPCA is a common technique in functional data analysis that decomposes the leading sources of variability among the curves 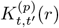 into a set of finite-dimensional, approximately uncorrelated principal components (PCs); see [36]. To do so, for each pair of cell types *t* and *t*′, we define the empirical covariance kernel as

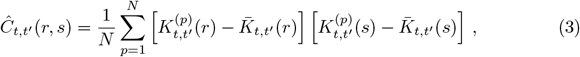

where 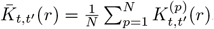.

The eigenvalues and eigenfunctions of the kernel *Ĉ*_*t,t*′_ are then computed to satisfy the functional equation

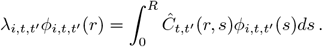

The *K* function 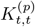 is summarized using the *d* coefficients (PCs)

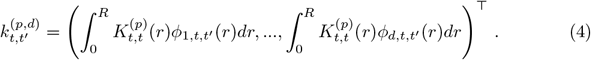

The coefficients comprising 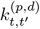 describe the projection of the *K* function 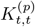 onto the finite dimensional linear space spanned by *ϕ*_1,*t,t*′_, …, *ϕ*_*d,t,t*′_, which are optimal in terms of capturing the variability among the curves 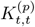, *p* = 1, …, *N*, with a *d*-dimensional summary. We note the amount of captured variability can be computed to give estimates for the effectiveness of the components in explaining the functions, and further used to determine the number of required components to achieve some level of fit. Another advantage of summarizing the curves in this way is that, when differences in the K functions across the population are present due to differences in the outcome(s) of interest, the PCs are expected to capture these differences.

As such, we summarize the spatial data using the principal components 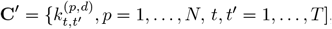, which we then incorporate with the meta-data **M** into a model for **Z** of the form

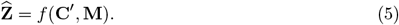

Since our ultimate goal includes evaluating which spatial interactions or elements of the meta-data are useful in predicting the outcomes, we use a random forest model for *f*. Random forest models are tree-based ensemble machine learning methods in which decision trees are built, after sampling with replacement the patient data and discarding some covariates at random, by sequentially splitting on variables to minimize a metric for predicting **Z** [54]. The main reasons for the sampling procedures for the patient data and covariates in building each tree is to build nearly independent trees and also address overfitting, common in many machine learning applications. Data missing from one tree is likely present in another tree. When the trees are combined to create a forest, increased statistical power is observed. When outcomes in **Z** take the values 0 or 1, as in the TNBC data, majority voting is used for the model estimates.

### Variable importance

Random forest models are useful in achieving our goals since they have strong predictive power while still allowing for a quantification of the usefulness of individual covariates in predicting the response through various “variable importance” measures. However, there are several challenges to overcome in calculating and quantifying the significance of such importance measures in this setting. One is that the variables in **C**′ are *d*-dimensional proxies of the information derived from the spatial image data. When multiple components are used to describe a single function, i.e. *d >* 1, we must take into account that each individual component in **C**′ describes only a portion of the associated *K* function. Therefore, the importance of each component must be combined to describe the importance of each spatial interaction, and further this importance must be made comparable to that of the meta-variables. Also, we wish to identify spatial interactions and meta-variables that are of “significant importance”, which we take to mean that their importance exceeds to a statistically significant degree that of similar variables that are unrelated to the response. This task is complicated by the fact that we are often faced with such a large number of spatial interactions. Given the large number of variables, we expect some to have anomalously large variable importance even when they are independent from the response.

Before proceeding we now describe the computation of variable importance metrics for random forest models, some of which were introduced in the seminal paper on random forests [54]. There are multiple methods for calculating variable importance values, such as permuting variables in data and comparing the difference in loss metrics, or measuring node purity. We tested several methods and found similar results regardless of the variable importance metric calculation. We implement a node purity metric, sometimes called recursive partitioning, as described in [55] and [56]. Our choice is due to simulation and computation considerations, but we encourage future research on the effect of different metric calculations.

In the following explanations for node purity variable importance, it is perhaps useful to imagine the scenario where the outcomes *Z*_*i*_ take the values 0 and 1. When each constituent decision tree is formed in producing the random forest model, nodes are split based on some impurity metric relating to the outcomes *Z*_*i*_ [55]. For a given node, *A*, node impurity is defined as

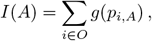

where *g* is an “impurity” function, *O* is the set of possible outcomes, e.g. 0 or 1, and *p*_*i,A*_ is the proportion of data in *A* which belong to outcome class *i* of *Z*, i.e. 0 or 1. Typical choices for *g* are the information index (*g*(*p*) = −*p* log(*p*)) or the Gini index (*g*(*p*) = − *p*(1 − *p*)). At a given stage of the decision tree, the splitting variable and location is chosen to maximize the impurity reduction

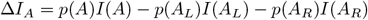

where *A*_*L*_ and *A*_*R*_ are respectively the left and right resulting nodes and *p*(*A*) is the probability of *A* (for future observations) [56]. That is, 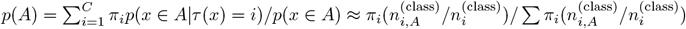 for *C* classes, prior probabilities *π*_*i*_, the classes assigned *C*, and the number of observations 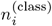 (that are class *i*) and 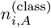 (that are class *i* in *A*). A variable *v*’s importance in a single tree can be computed as

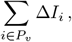

where *P*_*v*_ is the set of primary splits for (i.e. nodes that split on) the variable.

Variable importance (for the entire forest) can similarly be calculated by considering nodes across all decision trees, typically standardized by the number of trees fit or the number of trees where the split was present,

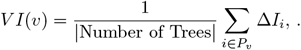

Additional modifications, such as the use of surrogates, can also be added to improve variable importance metrics. The technical details are left to more complete works, e.g. see [55]. The principle behind surrogate splits is that at each split a selection for the splitting variable must be made, even if the selection is between nearly identical variables. Traditional splitting would select only one, giving it a measure of variable importance, and ignore any other options. In this way, traditional splitting can result in masked variables and inaccurate representations of variable importances. A model using surrogate splits revisits each split after selecting the “primary” splitting variable and considers alternative, “secondary” splitting variables, mitigating the masking behavior.

Surrogate splits also take into account that the variables used after the primary split are, to some degree, lower quality and weight the splits accordingly. Weighting is performed according to a split’s impurity reduction, and varies slightly based on classification or regression problems [55]. Moreover, these surrogates allow for use of another split if an observation is missing data for the primary split.

Consider that a naïve choice for any variable missing data used in the primary split would be to go with the majority split. In order to evaluate potential splits, they should perform better than this naïve approach. In particular secondary splits should outperform this approach on missing data, and perhaps even split nearly as good as the primary split on all data. Consequently, one potential weighting scheme for node *A* and potential surrogate *v* is

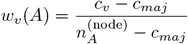

where *c*_*v*_ is the number of observations correctly classified due to splitting by variable *v, c*_*maj*_ is the number of observations correct by naïvely selecting the majority class, and 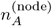 is the total number of observations in the node.

A typical variable may appear many times in a tree, both as a primary and surrogate splitting variable. Therefore, the overall measure of variable importance for a given variable is a combination of the splitting importances

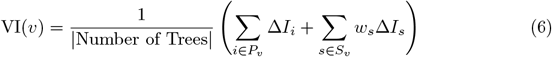

where for variable *v, P*_*v*_ and *S*_*v*_ respectively indicate the set of primary and surrogate splits.

### Variable importance comparison

Although the computed variable importance is a helpful summary statistic for ordering variables in terms of their expected usefulness in predicting the outcome, these values cannot be easily compared to determine which variables are significantly more useful than others, including those that are known to be independent of the response. Furthermore, after combining variable importance values from spatial variables in **C**′, these values must be compared with those from variables in **M**. Yet since each spatial interaction is characterized by *d*, which is typically larger than one, principal components, we might expect for example that the variable importance of a set of such components will exceed that of any meta-variable when both are independent of the response. Indeed, in totally random cases, with similar distributions, the spatial interaction term would often be selected approximately *d* times more than a meta-variable.

To allow for such comparisons, we standardize the variable importance metrics by adding matched synthetic variables. These synthetic variables are generated by permuting the true data in order to maintain identical distributions, but are independent from the outcome. We denote the synthetic spatial components as **C**′_*s*_ and the synthetic meta-variables **M**_*s*_.

)A random forest model is fit using both the true and synthetic variables,

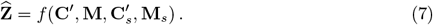

To build 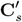, for each *b* iteration, 1 ≤ *b* ≤ *B*, where we take *B* = 100, a random functional variable is selected. The *d* principal components associated with each patient are permuted across patients, resulting in assignment of the *d* components to a random patient and hence outcome. Note that in doing so, the *d* components are kept together.

One could use *B* = 100 synthetic variables for each functional variable. However, investigations into the model through extensive simulations has shown that a single group of *B* synthetics is generally sufficient for the *K* functions, and additional synthetic *K* variables do not improve power beyond this.

Although one synthetic group for meta-variables could be used, previous work has shown a tendency for random forests to favor continuous predictors over discrete predictors [57]. The model accounts for this tendency through unique synthetic meta-variable groups. Therefore, **M**_*s*_ is created by permuting each meta-variable across patients *B* times.

We use these synthetic variables, 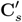 and **M**_*s*_, to standardize the variable importance values of the true data and build noise cut-offs. In doing so we are able to infer which spatial interactions and meta-variables lead to significant improvements in the model accuracy. The details of this are left to Comparing variable importance of spatial interactions and meta-variables to noise.

Due to the innate randomness in the models, the variable importance values fluctuate between runs and model fits of the random forests. To quantify this, we employ cross-validation (CV). CV begins by splitting the data into *F* roughly equal-sized parts, sometimes called folds. On each iteration one of these parts is left out, and the remaining data are used to fit the model. The fluctuating variable importance values between runs can be used to estimate the variability. Moreover, the fitted model can be used to predict the outcomes in the left out part/fold in order to evaluate the predictive power of the model.

Let the data be randomly assigned to the *F* folds such that an indexing function *κ*: {1, …, *N*} ↦ {1, …, *F*} indicates the partition to which each *p*^th^ patient’s data are allocated. Denote the fitted model, using the true variables as in model (7), with the *j*^th^ fold removed 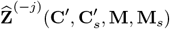. The estimated CV variable importance for each functional variable *c* (in both **C**′ and 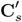), which are described by the previously discussed *d* dimensional principal components 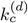, is computed as

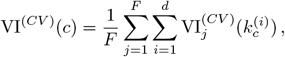

where 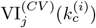 denotes the variable importance estimate from the *j*^th^ fold-removed model 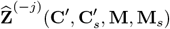 for component 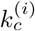, which is the *i*^th^ component relating to the *c*^th^ function.

The estimated CV variable importance effects for a meta-variable *m* (in **M** and **M**_*s*_) are similarly computed as the mean variable importance metric,

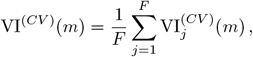

such that 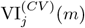 denotes the variable importance estimate from the model 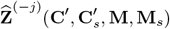 for meta-variable *m*.

We quantify the uncertainty in the estimate of the variable importance measure for each variable *v*, both functional and meta, by calculating its standard deviation across the *F* folds.

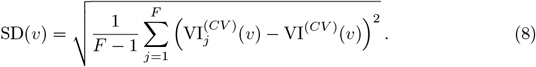

This uncertainty estimate is used with variable importance estimates created from the mean of non-cross-validated models. Let VI_*j*_(*x*) indicate the variable importance metric from model (7), iterate *j* (where we take 1 ≤ *j* ≤ *F* for ease, but each run is on the full data set), then the estimates are computed

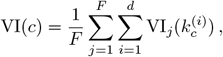

for functional variables, *c*, and

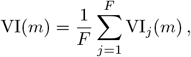

for meta-variables, *m*.

### Comparing variable importance of spatial interactions and meta-variables to noise

As mentioned above, due to the use of *d* ≥ 1 principal component summaries, we expect that for spatial interactions and meta variables that are independent of the outcome the variable importance of the spatial interactions will typically be larger. As such we use the estimated variable importance values for the synthetic variables to calibrate the variable importance between the spatial interactions and meta-variables. We compute for each of the spatial and meta-variables, respectively *c* and *m*, the empirical *α* quantiles of the variable importance values of the synthetic variables. If **C**^(*s*)^ indicates the set of synthetic functions, i.e. the combined synthetic components, and **M**^(*s,m*)^ indicates the set of synthetic meta-variables matched to meta-variable *m*, then we set

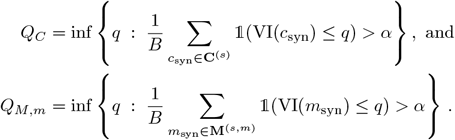

Let **Q**_*M*_ = (*Q*_*M*,1_, …, *Q*_*M*,|*M*|_) where |*M*| is the total number of meta-variables. Below we always set *α* = 0.95. Letting *Q*_noise_ = max{*Q*_*C*_, **Q**_*M*_}, we calibrate the variable importance of each true variable computed from model (7) that includes synthetic variables, denoted *v*_true,(7)_, as

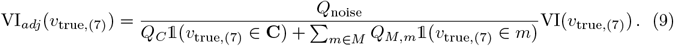

The cross-validated standard deviations of the variable importance values are similarly adjusted, but based on the model (5) that does not include the synthetic variables, denoted *v*_true,(5)_, as

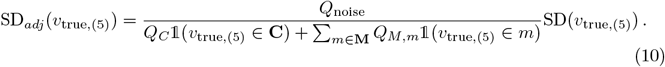

These adjusted variable importance values may then be compared to *Q*_noise_. Estimates that go below the cutoff *Q*_noise_ have variable importance values that appear to show no statistically significant relationship with the outcome. Plotting *Q*_noise_, the adjusted variable importance values, VI_*adj*_(*v*), along with their adjusted standard deviations gives a simple way to evaluate at a glance which variables appear to be important in predicting the response. An example plot of this is shown in the right hand panel of Fig 2, where *Q*_noise_ is plotted as a vertical, red-dotted line, and the variable importance values are plotted along with their adjusted standard deviations.

While *Q*_noise_ estimates the *α* percentile of the variable importance values corresponding to outcomes that are unrelated the outcome, the variable importance values themselves can be distorted due to overfitting. Overfitting occurs when the model fits well to the observed data but does not generalize well to unobserved data. As more variables are observed, it is increasingly likely to find a variable that seems to have high importance, despite no true relationship to the response.

To account for this potential effect, we employ an additional variable permutation. In this step, we permute the true variables in addition to the synthetic variables, *H* times. We take *H* = 100 below. In each forest, the synthetic variables are again used to align the variables (using the previously computed *Q*_noise_), but the remaining, aligned “true” variables do not have any relation to the outcome. In each of these forests based on variables that are independent from the response, 1 ≤ *b* ≤ *B*, the resulting, adjusted variable importance estimates are ordered,

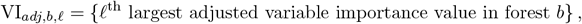

ensuring VI_*adj,b*,1_ *>* VI_*adj,b*,2_ *>* … *>* VI_*adj,b,V*_ where *V* denotes the total number of variables between the spatial interactions, **C**, and the meta-variables, **M**. When the number of variables *V* is large, especially in relation to the number of patients *N*, we might expect even when the spatial variables and meta-variables are independent of **Z** that the largest variable importance values will (far) exceed *Q*_noise_. As such, we also compute the *α* quantile of the variable importance values in each ordered position *ℓ* for the random forests fit to the permuted data,

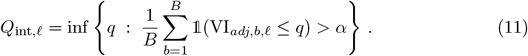

We let **Q**_int_ = (*Q*_int,1_, …, *Q*_int,*V*_), and coin it the interpolation cutoff.

We include **Q**_int_ with *Q*_noise_ in order to quantify overfitting and thereby evaluate the significance of values of the variable importance values VI_*adj*_(*v*); this is the orange-dotted line in the right hand panel of Fig 2.

In summary, variables with adjusted importance values, VI_*adj*_(*v*), that are larger than both *Q*_noise_ and *Q*_int,*ℓ*_ exhibit importance that significantly exceeds (at the 1 − *α* level) what we might expect from similar variables that are unrelated to the outcome. This holds taking into account the inflation in the variable importance values that arise from fitting the random forest to a large number of spatial interactions and meta-variables.

### Predictive accuracy estimates

In weighing the significance of the computed variable importance values, one should also consider the overall predictive accuracy of the final model for the outcomes. For example, a variable may have a large variable importance value within a model that does not lead to improved predictions of the outcome over naïve models. Therefore, evaluating the importance of a variable in the context of a random forest model requires some consideration for model accuracy.

We consider three such estimates of the overall predictive accuracy of the model: out-of-bag accuracy (OOB), naïve guess (GUESS), and biased guess (BIAS). OOB estimates how well the model works on unseen data, and a comparison of this estimate to GUESS or BIAS can indicate if the model captures additional information beyond simplistic estimates.

OOB estimates how well the model would perform on new data and is computed by predicting the data left out during each CV iteration,

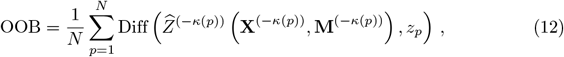

for the *N* patients and where Diff indicates a difference function. For classification problems such as with the TNBC or LUSC versus LUAD data, this may be defined using an indicator function,

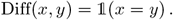

GUESS is computed by randomly guessing the outcome based on the outcome’s observed frequency in the original data. To account for the natural variability in this estimate, we use the mean estimate of *G* trials,

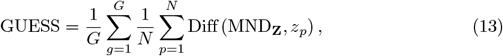

where MND is a multinomial distribution with probability of each outcome based on its observed frequency. For example, in the TNBC data in which we wish to predict the “compartmentalised” versus “mixed” result with a proportion *p* of “compartmentalised” patients, this amounts to computing the rate at which we would accurately guess the outcome by flipping a coin independently for each patient with probability *p* of heads, and guessing the outcome is “compartmentalised” for heads, and is “mixed” otherwise. For a random patient drawn from this two-outcome data, this approach will have a success probability of *p*^2^ + (1 − *p*)^2^. We take *G* = 500 below.

BIAS is built by always guessing the most likely outcome. For classification problems, the most likely outcome can naturally be defined by the data mode. We compute

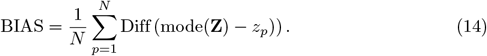

For two outcome data sets like the TNBC or lung cancer examples, this will have success probability for a random patient drawn from the sample of max{*p*, 1 − *p*}.

Similar naïve estimates can be derived for regression problems.

### Variable importance plot

The variable importance plot summarizes the variable importance values of both the spatial interactions and meta-variables. It shows how they compare to what we might expect from similar noise variables which are unrelated to the outcomes, and also summarizes the overall efficacy in predicting the outcomes using the random forest model. Fig 3 is the variable importance plot created from simulated data with two cell types, *A* and *B*, and two meta-variables with differing distributions, *age* and *sex*.

**Fig 3.**
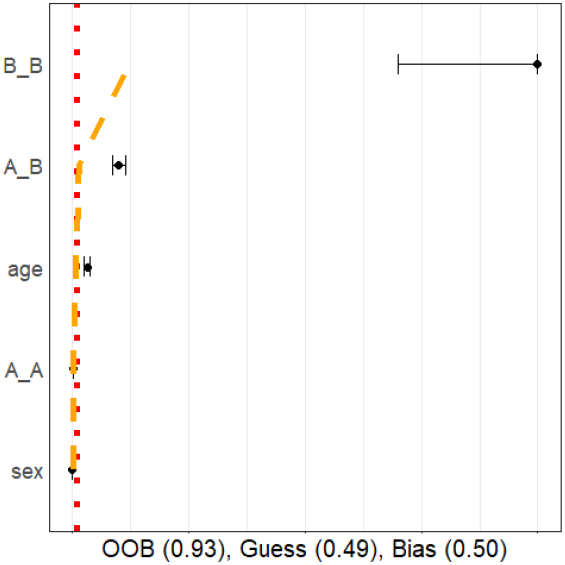
Sample variance importance plot. This sample variable importance plot uses simulated data with a binary outcome, two cell types, and two meta-variables. The data was simulated with significant differences between the outcomes for the in *B B, A B* spatial interactions, and age meta-variable, but no significant difference across sex and the *A A* spatial interaction.

Variables are displayed according to their adjusted, standardized (0, 1) variable importance estimates (black dots), with the largest values being on top.

The plot visualizes uncertainty in computing the VIs through (gray) intervals representing one standard deviation on either side of the estimate. These intervals are the adjusted standard deviations estimated through CV, as given in Eq (10).

The noise cutoff *Q*_noise_ as well as the interpolation cutoff **Q**_int_ at levels *α* = 95% are also shown. Fig 3 presents the noise cutoff as a red, dotted, vertical line and the interpolation cutoff as a orange, dashed, curved line.

Estimates for predictive accuracy are given at the bottom of the figure. The estimates include OOB, GUESS, and BIAS, respectively defined in Eqs (12), (13), and (14).

Further interpretation of the variable importance plots are given in the sections Simulation study and Applications to MIBIscope data. These sections consider simulations for which there is a known solution and real data examples that can be interpreted.

## Results

In order to obtain some confidence for the fit of the proposed model on real data, in this section we first present simulations then address the patterns of interest.

### Simulation study

We here demonstrate our methodology via simulations of spatial point patterns. We produced simulated point patterns with properties, such as cell counts, numbers of phenotypes, etc., similar to that of the TNBC data set in [12]. The real data motivating this simulation experiment are analysed in the following section.

We considered simulated data from 34 “patients”. These patients were defined as being negative or positive for a binary outcome *Z* (note that positive/negative here refers to the arbitrary outcome *Z*, and is not related to hormone receptor status as it does in the term TNBC). We let there be 17 positive and 17 negative patients and simulate one image per patient. Each image consists of a point pattern with 16 cell phenotypes. Patients were also ascribed a single meta-variable, which we call age. We developed a random forest model as described in Materials and methods to predict patient outcome using interactions between the 16 different cell types in the images and the additional meta-variable.

We considered two main settings: (1) a simulation with only non-informative variable-outcome relations and (2) a simulation with both informative and non-informative variable-outcome relations.

To generate the images, the cells were placed according to multiple, potentially nested, (modified) Thomas processes, which are constructed iteratively [58]. For a given image, a Thomas process first places cells (of a given type, say *c*1) at random, according to a Poisson process. The Poisson process selects the number of cells to place, *n*_*c*1_, randomly based on a Poisson distribution, where the distributional parameter is user selected. For our simulation study, these are selected to correspond to the mean number of cells in the TNBC dataset. We standardize the images to unit length in both the *x* and *y* directions.

Around each cell *c*_*a,c*1_, *a* ∈ {1 … *n*_*c*1_}, cells of a different type, say *c*2 may be placed. Again the number of cells of *c*2 are randomly selected based on a Poisson distribution with a user selected parameter. The coordinates of the *c*2 cells are placed according to another distribution in such a way that they either cluster or disperse around the *c*1 cells. In our experiments, a bivariate Normal distribution is used to place the *c*2 points, so that the mean coincides with the location of a randomly selected *c*1 cell, and the covariance matrix is a scalar multiple *σ*^2^ times the identity matrix. By varying *σ*^2^ in the outcome groups, clustering or repulsion in the interactions between the cells can be introduced.

Additional cells can be simulated around the points of *c*1 or *c*2, and so on. Moreover, the original cells can be removed such that the new cells exhibit hidden or self-clustering. Compilations of such placement patterns, with use of potentially different distributions, can achieve images with varying degrees of clustering or regularity.

In our simulations, 16 cells were iteratively placed according to this modified Thomas pattern. Some cells were placed completely at random (*c*1, *c*4, *c*5, *c*6, *c*7, *c*13, *c*14, *c*15), some were placed exhibiting self-clustering (*c*8, *c*9, *c*10, *c*11, *c*12), and some were placed exhibiting clustering around *c*1 (*c*2, *c*3, *c*16). In what we call the “no–relation” simulation, the cells locations were simulated in the same way for both the positive and negative patients. However in the “relation” simulation, *c*2 exhibited increased clustering around *c*1 while *c*3 exhibited repulsion from *c*1, for positive outcomes. Similar to the true data, some cell types were present hundreds of times per image while others only rarely appeared. Each of the synthetic cell types were generated to mimic behaviors and frequencies seen in the TNBC data. Fig 4 presents two images, an image from the TNBC data set and a simulated image.

**Fig 4.**
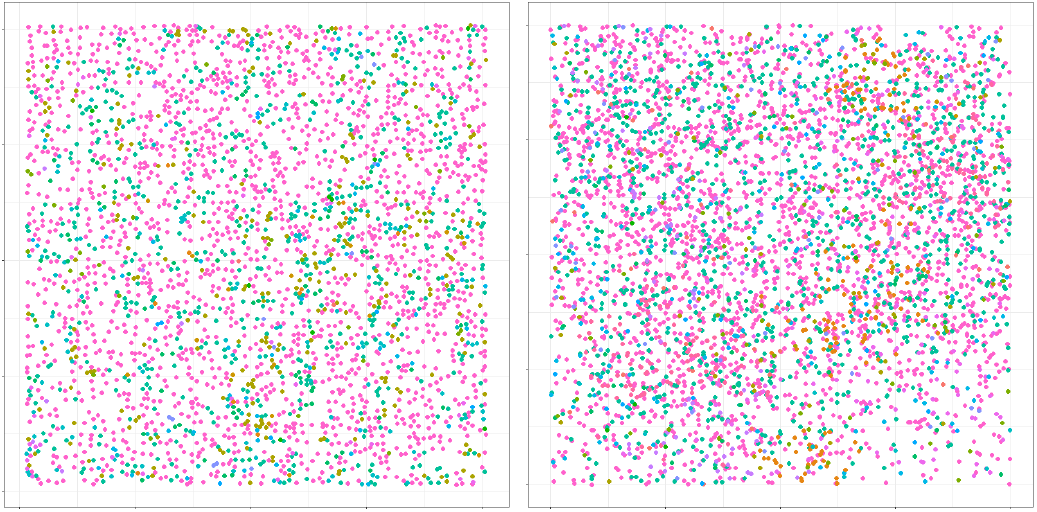
Comparison of TNBC and simulated data. An image from TNBC data (left) and simulated data (right). Different colors indicate one of the 16 different cell phenotypes.

In this way a single image for each of the 17 positive and 17 negative patients were simulated. The meta-variable age was simulated as a Normal random variable with unit variance. While in the no-relation case the mean was a constant 25 for both outcomes, in the relation-case the mean of age was set to be 25 for negative outcome patients, and 27 for positive outcome patients, creating a useful, though imperfect, classifier.

Moreover, in the relation-case, 2 of the 17 positive patients were given no-relation to the outcome as additional “noisy” images.

When modeling both cases, there were several tuning parameters selected. We used the standard choices of 500 trees for each random forest, 10 folds in cross-validation, and a standard significance level of *α* = 0.05. We used 100 interactions for the permuted random forest in creation of the interpolation cut-off. In (unreported) numerical investigations of these values we saw robustness in the results for these choices. Creation of the *K* functions also required selection of the maximum radius, *R*. Although we investigated the effect of the choice, we saw little variation in the results and used a traditionally recommended 25% of the side length of the image, along with the previously discussed isotropic edge corrections for the simulations. Moreover, the *K* functions were summarized using 3 principal components, which were selected to match the TNBC analysis. TNBC uses 3 components to explain at least 95% of the total variance explanation for each *K* function.

Fig 5 shows representative variable importance for all variables (left) and the top 25 (right) in the “no–relation” case. One may see from the plot that some variable importance estimates exceeded the red noise cutoff, but they all were observed to be below the orange interpolation cutoff. This may be interpreted that the observed variable importance values did not exceed, to a significant degree, what we would expect to see from similar variables that are known to be independent of the response.

**Fig 5.**
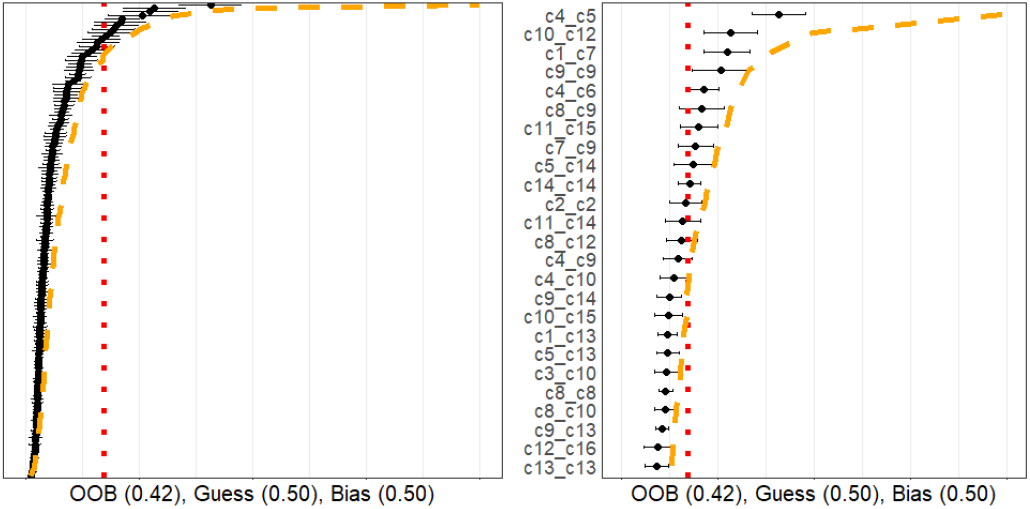
No relationship simulation. Simulation of 16 cell types for 34 patients with meta-variable age. The figure on the left gives the variable importance values for all variables while the one on the right shows only the top 25. All variables were generated with no-relationship to the outcome.

Moreover, the prediction accuracy estimate (OOB) indicates the model performs similarly to a naïve guessing approach. Taken together, the plot indicates that none of the spatial interactions appeared to be important in predicting the outcome, as was expected in this case.

On the other hand, Fig 6 shows a representative variable importance plot computed from a single simulation run in the “relation” simulation. All variables are shown in the left hand panel, and only top 25 are shown in the right panel. Although many variables are still below the noise and interpolation cutoffs, the known related variables are found to have a significant relationship with the outcome variable. Moreover, the OOB estimate far exceeds that of the GUESS or BIAS estimates. This plot indicates that interactions between the *c*1, *c*2, *c*3 cell types, and the age variable, appeared to be useful, to a statistically significant degree, in predicting the outcome, once again as expected. We note that since *c*2 and *c*3 cells are distributed around *c*1 cells, any change in these distributions will necessarily lead to differences in the cross *K* functions between these two cell types as well, as observed.

**Fig 6.**
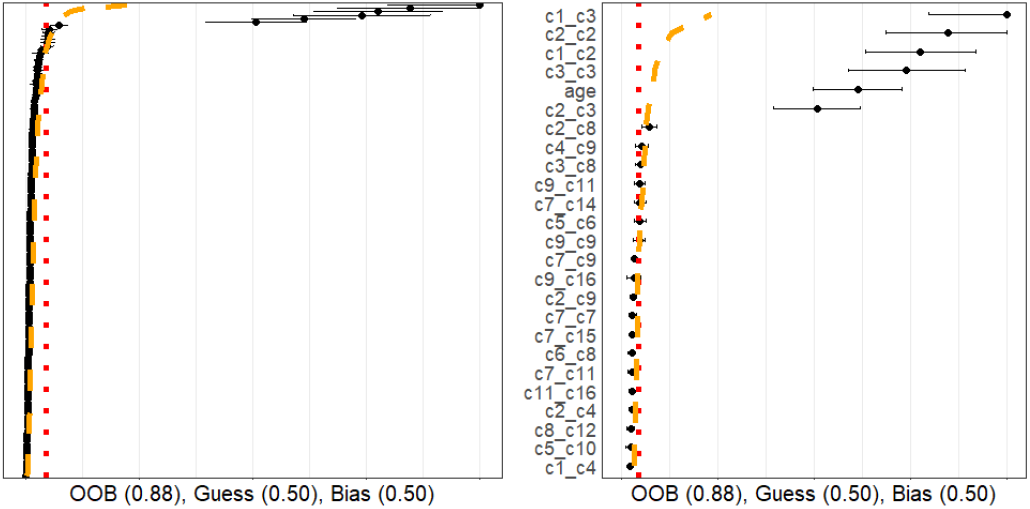
Relationship simulation. Simulation of 16 cell types for 34 patients with meta-variable age. All variable importance values (left) and top 25 (right) are shown. Most cell types were generated with no relationship to the outcome. However, age, *c*1 *c*2, and *c*1 *c*3 were designed to have a relationship with the outcome (which naturally means *c*2 *c*2 and *c*2 *c*3 would also have relationships to the outcomes). These variables are seen with significantly larger variable importance values.

In order to verify that each of the orange and red lines appeared to be appropriately calibrated, we performed an additional simulation experiment. We considered two cases (1) with only 4 cell types and (2) with 16 cell types. Each cell type other than cells *c*1 and *c*2 are generated such that they have no relationship to outcome. We modify the clustering of *c*2 around *c*1 in the positive group, and examine the rate at which the variable importance estimated for the *c*1 *c*2 spatial interaction exceeded the various cutoffs (red/orange lines). By changing the standard deviation in the Thomas process for placing *c*2 points around *c*1 points, we were able to investigate whether the approach is able to detect the presence of a relationship when the cells either cluster or are more dispersed across the binary outcomes. The resulting power curves, based on 100 simulations for each setting, are shown in Fig 7. These show the rate at which the variable importance estimates exceed the 95% noise cutoff, interpolation cutoff, both the noise and interpolation cutoffs, and the largest interpolation cutoff among all variables. In reference to Fig 7, the standard deviation in the Thomas process relating c1 and c2 takes the value of approximately 0.025 in the “no relation” case. As such, for this value it is desired that the noise and interpolation cutoffs are exceeded no more than *α* = 0.05 proportion of the time. Otherwise, detecting that standard deviation values smaller/larger than 0.025 lead to increased/reduced clustering of *c*1 *c*2 in the positive groups is desired.

**Fig 7.**
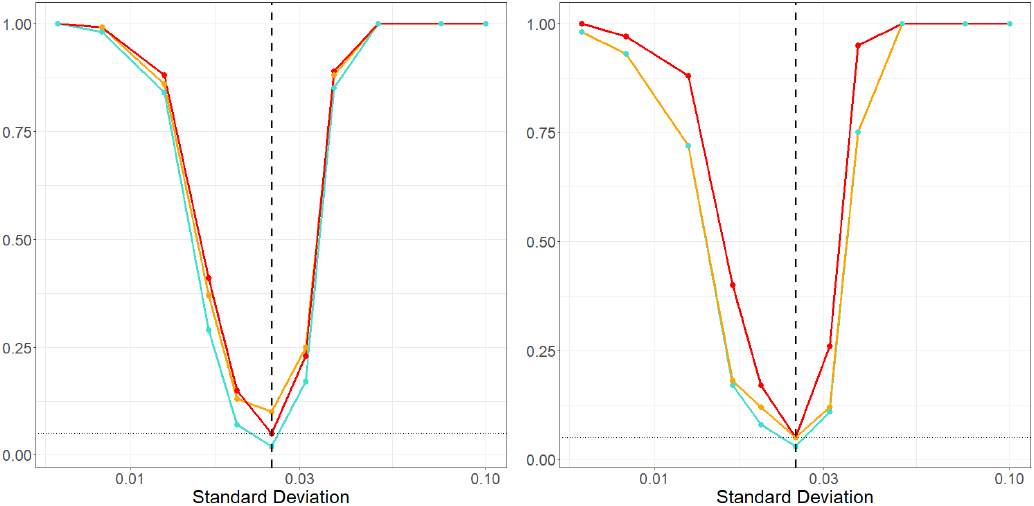
Power curves. Power curves showing the empirical rate, from 100 simulations in each setting, that the variable importance for the spatial interaction *c*1 *c*2 exceeded the 95% noise cutoff, interpolation cut off and both the noise and interpolation cutoff. The left image is based on spatial data with only 4 cell types and the right is based on 16 cell types. Colors match the variable importance plot when possible and indicate the cutoff method: above both cutoffs (teal), above the curved interpolation cutoff (orange), and straight noise cutoff (red).

We saw that each cutoff appeared to have good power to detect clustering or dispersion relationships. The noise cutoff was well sized, and the interpolation bound was slightly oversized. We saw though that comparing to both the noise and interpolation cutoffs (red and orange lines) yielded a slightly under sized cutoff with good power.

### Applications to MIBIscope data

In this section we present two applications of the proposed methods to MIBIscope data sets. The first investigates known clusters in tumors related to triple negative breast cancer tumours, while the second investigates unknown relations in tumors related to lung cancer.

#### Triple negative breast cancer

We investigate the TNBC data set in [12]. [12] employ a mixing score to categorise tumors based on their TME. The mixing score they use was defined as the proportion of immune cells touching tumor cells, and was calculated as the number of immune-tumor interactions divided by the number of immune-immune interactions for an image. They separate tumors into “compartmentalised” and “mixed” groups (and a “cold” group that we ignore), such that compartmentalised tumors tend to have tumor cells aggregated together with immune cells located around or away from the tumor cells, and mixed tumors tend to have tumor cells and immune cells mixed together. This makes a useful test data set on which to employ our method, as it provides two tumor groups that explicitly have different TMEs. The data set contains 33 patients with a single image per patient, 18 mixed tumors and 15 compartmentalised tumors. We define the outcome *Z* as 0 for mixed and 1 for compartmentalised, and we wish to predict the outcome using interactions between 16 cell phenotypes in the images and an additional meta-variable, age.

One tuning parameter with *K* functions is the maximum radius *R*. For this data we investigated using several options–25, 50, 100, 500, and 1000 micrometers. In all cases the conclusions were comparable, illustrating a robustness to the choice. The only (expected) difference is that as *R* increases, additional principle components, *d*, are required to capture the variations in the functions. With this observation and domain knowledge, *K* functions up to a maximum radius of 50 micrometers were used. The results of this analysis are shown in Fig 8. As expected, many interactions are shown to be non-significant. However, there are a reasonable number above both cutoffs and the predictive accuracy estimates agree that there are some important relationships in the data. Specifically, tumor cell-tumor cell interactions came as the top interaction, consistent with the characteristics of the “compartimentalised” tumors where little immune cells infiltrate the tumors and tumor cells are densely packed.

**Fig 8.**
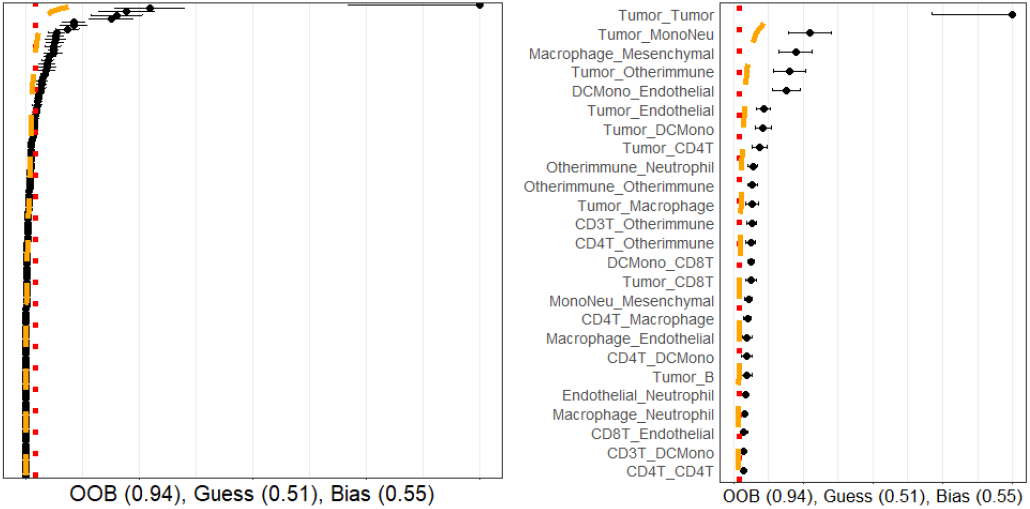
TNBC variable importance. Variable importance plot and random forest model summary for predicting “compartmentalized” versus “mixed” tumor types with the TNBC data. The OOB far exceeds those of naïve models, and many of the spatial interactions between tumor cells and immune cell populations exhibited significant variable importance values.

Although this analysis can indicate important relationships, it does not quantify the type of differences. While meta-variables can be easily compared using traditional statistical methods, the *K* functions are more difficult to analyze. To this end, we can also consider plots of the *K* functions. Fig 9 examines the Tumor Tumor interaction, which is found to be significant, and the CD4T Endothelial interaction, which was found to be insignificant. The significant *K* functions seem to be well separable and smooth (a property indicative of many cells being present in each image), while the insignificant *K* functions are not easily separable and have high variability in individual *K* functions of the same group.

**Fig 9.**
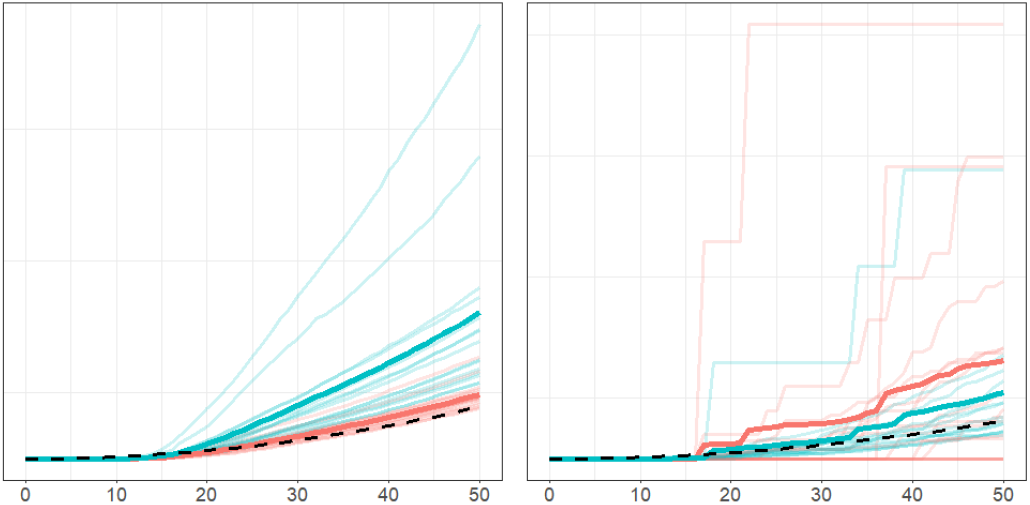
Example TNBC *K* functions. The *K* functions from the different outcomes compared. In both plots, the *x*-axis is radial distance in micrometers and the *y*-axis is the value of the *K* function. The light lines are the *K* functions for each patient, while the bold line indicates the average. In the figures, red indicates the mixed tumors while blue indicates the compartmentalized tumors. The black dashed line indicates the curve of a totally spatially random process for reference. The left figure is the Tumor Tumor interaction, which was found to have significant differences in the outcomes and the right figure is the CD4T Endothelial interaction, which was found have no significant differences between the outcomes. As expected, the compartmentalized group indicates increased clustering compared to the mixed group in Tumor Tumor cells while the CD4T Endothelial interaction shows no clear differences and varying *K* function patterns even within outcome groups.

#### Lung adenocarcinoma versus lung squamous cell carcinoma

We also applied our approach to attempt to predict different pathological subtypes of non-small cell lung cancer. In this section, we aim to measure to what degree the TME of two of the most common subtypes of lung cancer, LUAD and LUSC can be differentiated using the spatial relationships between phenotyped cells as characterised by *K* functions. Our dataset contained 44 LUAD and 20 LUSC samples stained with antibodies against 35 proteins to enable the phenotyping of tumor cells, fibroblasts, and 10 immune cell subsets after scanning on the MIBIScope. A summary of this data is given in S1 Table. *K* functions were computed between each cell type with again a maximum radius of 50 micrometers.

The variable importance plot and model summary when using our method to predict LUAD versus LUSC are shown in Fig 10. The results suggest that no spatial interactions as encoded by the *K* functions are significantly useful in distinguishing LUAD versus LUSC. The OOB accuracy was observed to be on-par with a naïve method, and none of the variable importance measurements exceeded the 95% quantile of what we would expect from independent point patterns. This indicates our method using homogeneous *K* functions applied to these specific cell phenotypes is unable to differentiate between the TME of LUAD and LUSC cancer types, and studying the data with additional cell subsets and/or through other summary functions or techniques may be of value.

**Fig 10.**
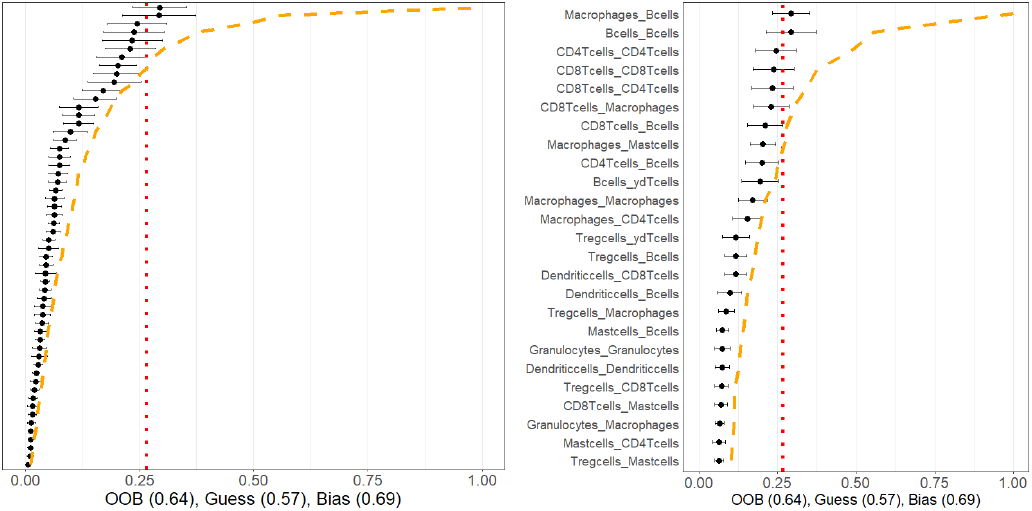
Lung cancer variable importances. Variable importance plot and random forest model summary for predicting LUAD versus LUSC tumor types. The OOB is similar to a naïve model, and none of the measured variable importance value were statistically significant.

## Discussion

In this paper, we present a novel method for the analysis of data-rich TME spatial data. We consider the case in which we compare TME and meta data or clinical data from two different patient groups, and develop a model to identify significant differences between these groups in the distribution of cell phenotypes, protein antigens, or a mixture thereof. In addition, the model aims to predict which group a new patient is in using their TME and meta data. Our model employs a combination of spatial statistics and functional data analysis and is applicable to marked point processes in general.

Benefits of the model include general applicability, with few tuning parameters, and easily visualised and interpreted output. We find our model to be robust to the choice of the tuning parameters. The model demonstrates a powerful ability to identify important variables while maintaining good predictive power.

We evaluate our approach on simulated data, demonstrating the effectiveness of our method for marked spatial point patterns with known interactions. Our approach is then applied to TNBC and lung cancer data obtained from multiplexed ion beam imaging. For the TNBC data, we compare two groups of tumors that are separated into “compartmentalised” and “mixed” groups based on the degree of tumor-immune cell interactions. This separation of tumors gives two groups with explicit differences in their TMEs, making a useful data set for the demonstration of our model. The model demonstrates good predictive accuracy, and identifies expected specific cell phenotypes relationships, and interactions of interest. The lung cancer data shows our model can also detect lack of differences in cell phenotypes relationships.

Throughout this paper we have assumed homogeneity of the underlying spatial point processes in defining the *K* functions used in the model. Spatial homogeneity is defined such that the intensity of a given mark is independent of spatial location [14–16].

Whilst such an assumption may be reasonable in some cases, given the complexity of the TME, homogeneity may not always apply. We note, however, that since our approach compares *K* functions between different groups, rather than against the theoretical *K* functions associated with complete spatial randomness (as is typical in other circumstances), the lack of underlying spatial homogeneity in the TME for a data set may not be overly problematic. Statistical tests for homogeneity exist [14–16].

Inhomogenous *K* functions can be used in an attempt to mitigate the issue of inhomogeneity [14–16, 59]. Regardless of how *K* functions are defined they are amenable to being used in our methodology. Furthermore, whilst we have focused throughout this paper on *K* functions, we note that the methods presented here can be applied to any summary functions from spatial statistics, or indeed any functions in general. See, for example, suggestions in [24, 60, 61].

In addition, we note that 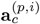 in Eq (1), may consist of the raw measured protein expression level for each protein, and may also include other cell information (e.g. cell size). Our approach could potentially be adapted for analyzing raw protein expression levels via the use of mark-weighted *K* functions [14, 62]. Consequently, the approach may be useful in methods applied to cell phenotypes based on proteins (OPAL, MIBI, Phenocycler Fusion) or transcripts (Xenium, Cosmx, MERscope) expression [39–45].

We have also assumed in this paper outcomes which are categorical, or classes in a group. In this way, we say the model performed classification. Although we considered two classes, a larger number of classes is directly possible. Further extension of this method to real-valued data, e.g. survival time, is likewise natural. Random forest models designed for continuous or survival time analysis exist, e.g. [54, 63, 64], and metrics such as *L*_2_ error can be used in place of the difference function of the OOB and naïve estimates.

## Supporting information

**S1 Table. Lung Cancer Summary Table**. Table summarizing the patients, number of images, type of lung cancer, and the cell count for each type.

